# Spatial Metabolomics co-registered with Multiplex Phenotyping for the Evaluation of Human Kidney Tissue

**DOI:** 10.1101/2023.11.10.566529

**Authors:** Greice M Zickuhr, In Hwa Um, Alexander Laird, David J Harrison, Alison L Dickson

**Affiliations:** School of Medicine, University of St Andrews, North Haugh, St Andrews, KY16 9TF; Department of Urology, Western General Hospital, Crewe Road South, Edinburgh EH4 2XU, UK

**Keywords:** Mass Spectrometry Imaging, DESI-MSI, multiplex immunofluorescence, image analysis, multi-modal imaging, AI

## Abstract

A workflow has been evaluated that utilises a single tissue section to obtain spatially co-registered, molecular, and phenotypical information suitable for AI-enabled analysis. The impact of varying DESI-MSI conditions (e.g., temperature, scan rate, acquisition time) on the detection of small molecules and on tissue quality for integration into typical clinical pathology workflows assessed in human kidney.

## INTRODUCTION

Spatial metabolomic technologies are crucial for understanding biochemical processes involved in the pathophysiology of diseases by capturing the metabolic heterogeneity of cell types within their tissue environment as metabolites are direct markers of biochemical activities, closely related to cell phenotypes^1,2^. Mass spectrometry imaging (MSI)-based metabolomics methods can map the distribution of hundreds of these chemical species and when coupled to other imaging modalities, MSI becomes a powerful technology. Conventional pathology techniques such as haematoxylin and eosin (H&E) stain, immunohistochemistry (IHC) and immunofluorescence (IF) provides complementary information on the spatial distribution of morphological features and specific protein markers.^3^ Data integration from the different imaging modalities is crucial to understand biological mechanisms.

DESI-MSI requires minimal sample preparation and is regarded as a “soft” ionisation technology as its non-destructive nature on tissue has been demonstrated using lab-built Sprayer^6^. Since introduced, lab-custom and the commercial Prosolia design have been used to study different pathologies and shown to be compatible with different imaging modalities (e.g. H&E, IHC, IF and IMC)^7–9^ allowing the obtention of phenotypical information from the same section submitted to MSI. These approaches, however, are limited to MSI followed by one other phenotyping technique. Furthermore, recently a new commercial sprayer and heated transfer line (HTL) were introduced, however, the impact of these recent technological improvements has not yet been assessed on tissue in a pathology or clinical setting.^10^

In this study case we report a workflow that utilises a single tissue section to obtain high molecular and phenotypical information suitable for AI-enabled image analysis. We assessed the impact of varying DESI-MSI conditions on the quality of tissue for integration on typical clinical pathology workflows and imaging research.

## METHODS

### Materials and Reagents

Polyvinylpyrrolidone (PVP) (MW 360 kDa), (Hydroxypropyl) methyl cellulose (HPMC) (viscosity 40−60 cP and 2600-5600 cP), methanol, water, iso-pentane, 2-propanol were of analytical grade or higher and purchased from Sigma-Aldrich (Steinheim, Germany). Haematoxylin (Mayers, pfm medical, UK), eosin Y 1% alcoholic and xylene (CellPath, UK), periodic-acid (TCS Biosciences Ltd, UK) and Schiff’s reagent (Merck, Germany). Description of the IF and IHC antibodies used can be found in the Supporting Information.

### Samples

Snap frozen non-cancerous kidney tissue was selected from partial nephrectomy undertaken as curative treatment of kidney cancer. Ethical approval was granted by Lothian Biorepository (SR1787 10/ES/0061).

### Tissue Embedding and Sectioning

Tissue was embedded in PVP (2.5%) modified HPMC (7.5%, 40-60 cP) which has been demonstrated previously by Dannhorn and colleagues to be compatible with tissue analysis^13^. Sectioning was performed on a dedicated (MSI use only) HM525 NX Cryostat (Epredia, Portsmouth, USA) to a thickness of 10 µm. Only mass spectrometry-compatible reagents were used in the cryostat to minimise the risk of source contamination. Serial sections were thaw-mounted onto Superfrost microscope slides (Thermo Scientific Waltham, MA), nitrogen dried, vacuum packed and stored in −80 °C. Prior to analysis sections were left to equilibrate to room temperature under vacuum for 20 min.

### DESI-MSI Experiments

Analysis was performed on a Xevo G2-XS Q-ToF equipped with a DESI-XS ion source and heated transfer line (Waters, Milford USA) operated in negative ionisation mode between *m/z* 50-1200. A solvent mixture of 98% methanol and 2% water was delivered at 2 µL/min and nebulised with nitrogen at a backpressure of 1 bar. Spatial resolution was set at 20 x 20 µm. Imaging experiments were performed by setting the transfer line temperature to low 150 °C and high 450 °C, and scan speeds of 10, 20 and 30 scans/sec.

Data processing and visualization were performed in HDI® (Waters, Milford USA) and normalised to TIC. Tentative compound assignments were made with high mass accuracy measurements (≤ 5 ppm mass error) using the Human Metabolome Database and Lipid Maps®. Tissue segmentation (UMAP) was performed using the Waters MicroApp MSI Segmentation, version 2.1.1 (Waters, Milford USA).

### Histochemical staining, multiplex immunofluorescence and multiplex immunohistochemical labelling and imaging

Following DESI-MSI, tissues were histochemically stained by haematoxylin and eosin (H&E) and/or periodic acid Schiff (PAS). An immunofluorescence panel containing NucBlue™ for nuclear identification and antibodies against p57 (glomerular podocytes) and PFKFB3 (marker of glycolytic activity) was optimised separately and applied to sections following DESI-MSI and histochemical staining (Supporting Information Table 1). Fluorophores were stripped off the tissues and a mIHC panel of vimentin and pan-cytokeratin was applied. An additional optimised panel for CD45^+^ CD8^+^ mIF for confirmation of lymphocytes was applied to cases of immune infiltration. For detailed method information see the Supporting Information. Brightfield images of H&E, PAS and mIHC stains and fluorescence images of mIF were acquired using a Zeiss Axio Scan Z1 scanner. A uniform scanning profile was used for each fluorescence channel. QuPath was used to visualize and export high resolution brightfield images.

### AI-enabled Image Analysis

Brightfield and fluorescent images were analysed using HALO® v3.6.4134.166 and HALO® AI v3.6.4134 (Indica Labs). For mIF image analysis a customized nuclear segmentation classifier was built using the NucBlue™ channel as a guide to negate falsely segmented nuclei^14^. Fluorescence intensity thresholds were set individually to each channel using the nuclear marker as a guide.

## RESULTS

A reproducible workflow for metabolomics analysis by DESI-MSI followed by histology and protein labelling on the same tissue section was optimised and evaluated (Figure 1).

**Figure 1.**
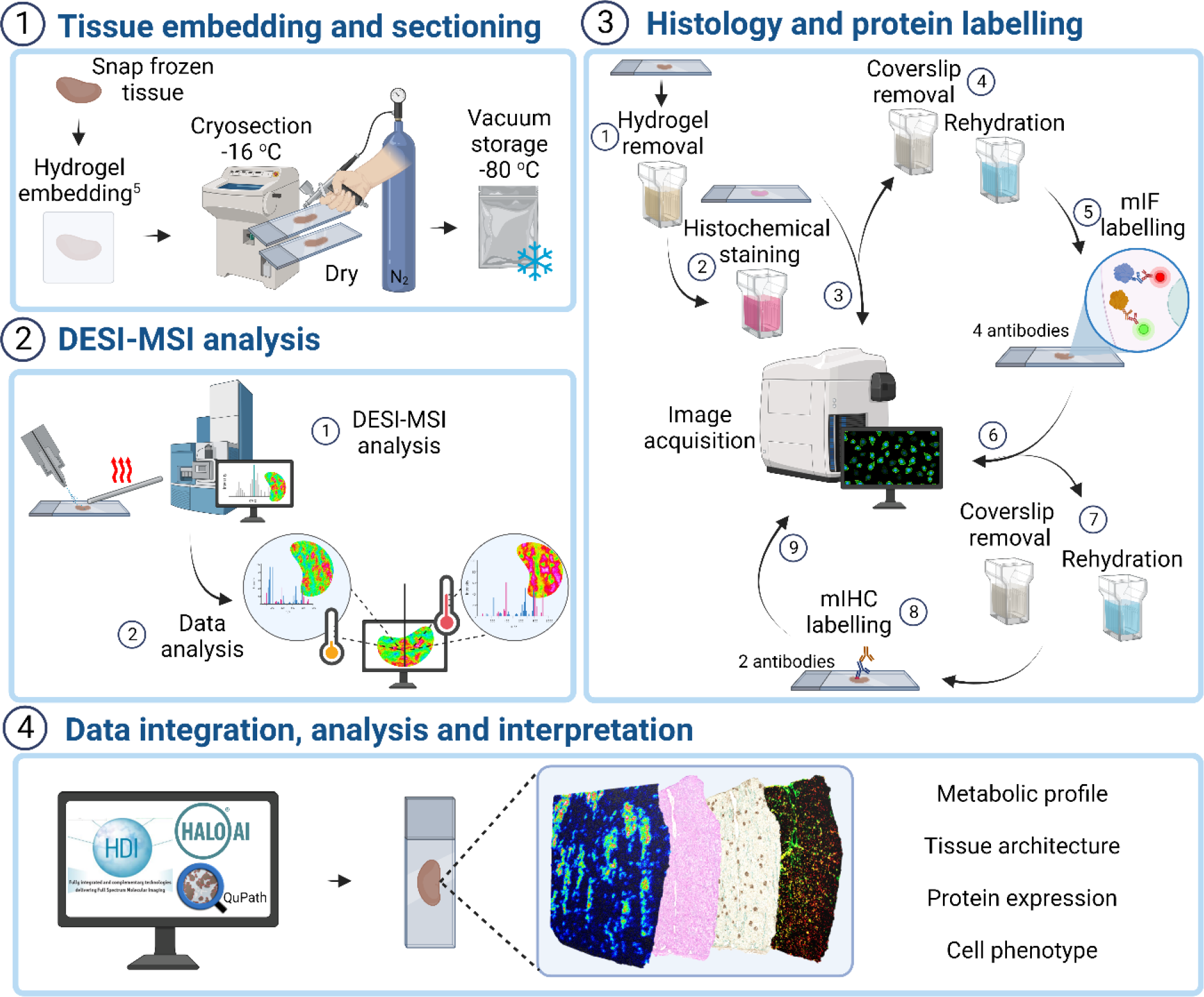
Schematic representation overview of the single-section multimodal imaging analysis pipeline highlighting key steps of **(1)** tissue section preparation for DESI-MSI; **(2)** tissue analysis by DESI XS MS with the redesigned Waters™ high-performance (HP) sprayer and heated transfer line (HTL); **(3)** post-DESI-MSI histochemical staining, multiplex immunofluorescence (mIF) and immunohistochemistry (mIHC) labelling steps and image acquisition and **(4)** analysis, integration and interpretation of data using HDI™, HALO® AI and QuPath for spatial correlation of metabolic profile and cell phenotype.

### Effect of heated transfer-line temperature and scan rate on compound classes detected by DESI-MSI

To determine the effect of the heated transfer line (HTL) on the signal intensity of different molecular classes, kidney tissue sections were analysed in triplicate with the HTL temperature set at 150 °C and 450 °C. The 1000 most intense *m/z* values from across all replicates generated a target list of 2455 features. A HTL temperature of 450 °C resulted in an increase of 1.8-fold in the overall signal intensity of species between *m/z* 600 to 1000 (e.g. lipids) (Figure 2A). In this range, 55.0% (628 out of 1142) of the features had a boost in signal greater than 1.5-fold and 38.4% greater than 2-fold. Features presenting the highest increase in signal were distributed within the tissue, whilst the most significant drops in intensity were in background peaks (fold-change (FC) 450/150 < 0.09). Lipids were putatively identified based on their exact mass, and their distribution to kidney features was co-registered using H&E images (Supplementary data). At the low *m/z* range (*m/z* 50-350), 488 peaks were detected, of which 52.3% had an average signal increase greater than 1.5-fold at 150 °C. Among these we identified lactate (*m/z* 89.0246, FC 3.22), succinate (*m/z* 117.0193, FC 3.86), glutamate (*m/z* 146.0458, FC 2.25) and arachidonic acid (*m/z* 303.2330, FC 2.29). Intense *m/z* signals arise in this mass region due to the chemical properties of the microscope slides; this was also affected by the lowest transfer line temperature (e.g. *m/z* 255.2332 and 283.2645). Interestingly, the intensity of taurine (*m/z* 124.0074, Figure 2B) was 2-fold higher at 450 °C whilst hypoxanthine (*m/z* 135.0311, FC 1.19) and ascorbic acid (*m/z* 175.0247, FC 0.78) were not highly influenced by the change in temperature. Figure 2C shows the effect of the sensitivity increase on the heatmaps of selected ion species.

**Figure 2.**
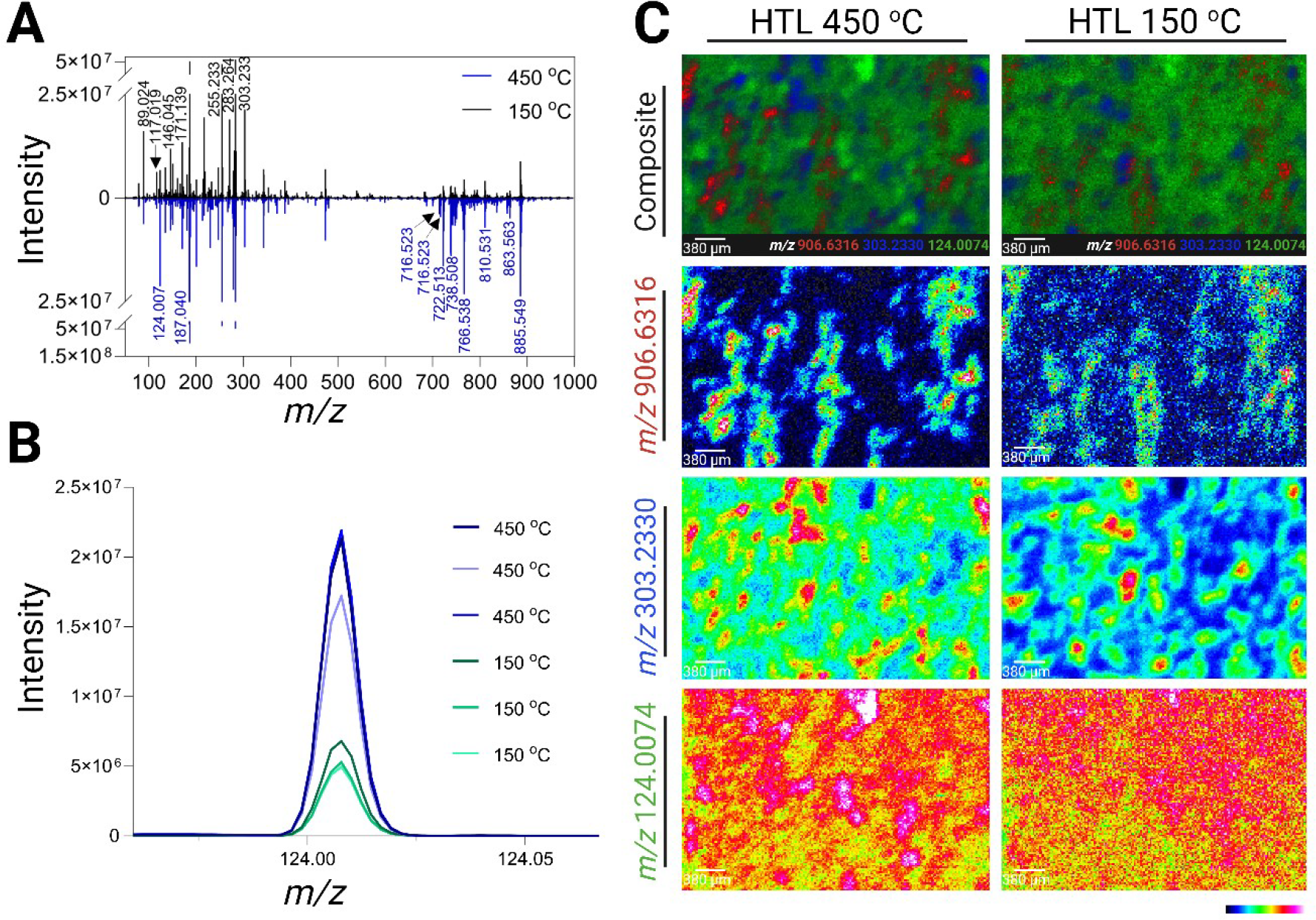
Effect of the DESI heated transfer line (HTL) temperature at 150 °C and 450 °C on different molecular classes of kidney tissue. Differences in **(A)** the negative ionization mode mass spectra of a kidney tissue analysed with HTL at 150 °C (black) and 450 °C (blue); and **(B)** mass spectrum of *m/z* 124.0074 (taurine) showing higher intensity with HTL at 450 °C (n=3). **(C)** DESI ion images showing better resolution of *m/z* 906.6316 (SHexCer 42:1;O3) and *m/z* 124.0074 (taurine) at 450 °C, *m/z* 303.2330 (arachidonic acid) at 150 °C and composite image of the three ions at both HTL temperatures.

**Figure 2.**
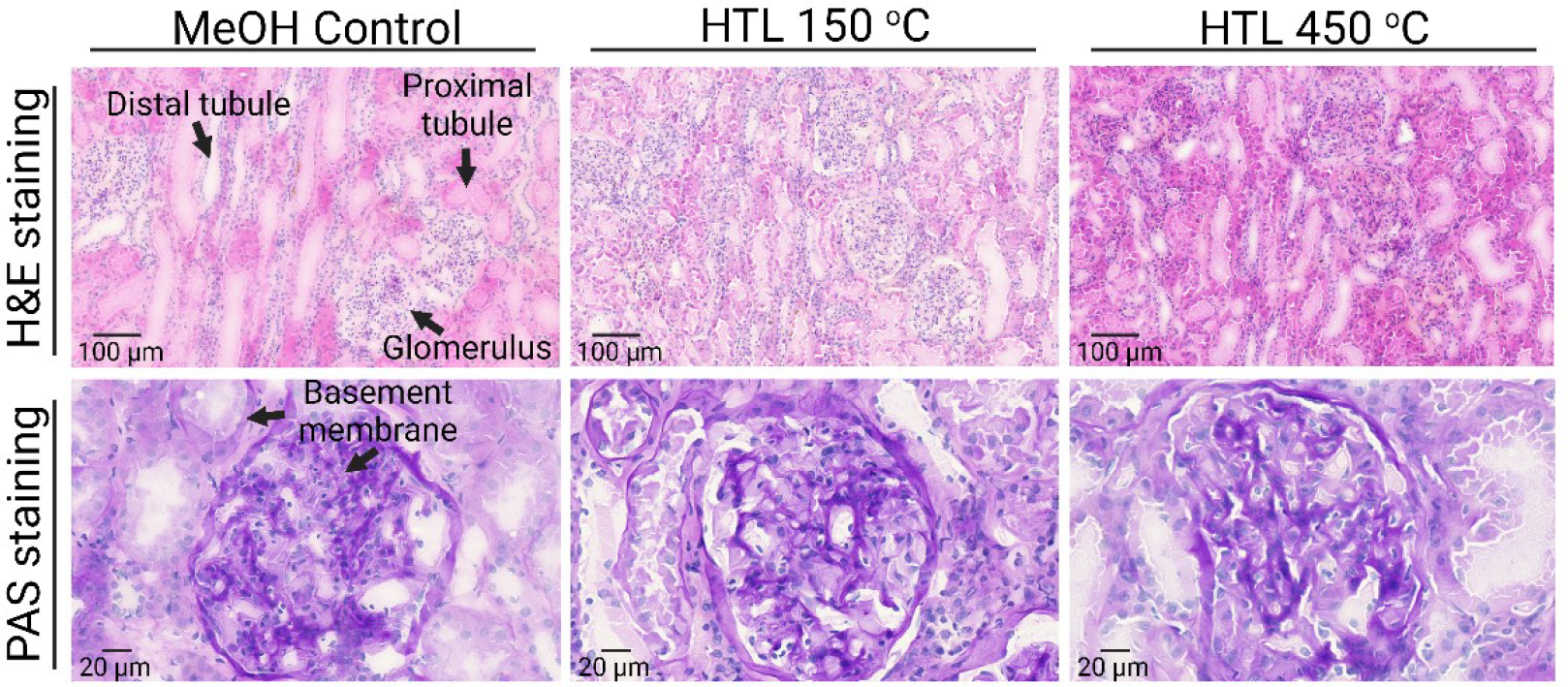
Histopathology assessment of kidney sections post-DESI-MSI. Haematoxylin and eosin (H&E) stain of MeOH control and post-DESI analysed tissue highlighting protein fixation as an effect of the interaction of solvent with tissue proteins and intensified by the higher temperature from the heated transfer line (HTL). Periodic acid Schiff’s (PAS) staining on the same sections highlighting intact basement membranes post-DESI-MSI submitted to both HTL temperatures.

As a HTL temperature of 450 °C was shown to result in higher signal for the lipid range we decided to evaluate the effect of different scan speeds at this temperature. The acquisition speed was set to 10, 20 and 30 scans/sec, with a fixed 20 x 20 µm pixel size, and resulted in average run times of 368, 216 and 143 min, respectively. A total of 6044 pixels approximately in the same area of the different scanned sections were extracted and raw data examined. Although signal intensity reduced with the increase in scans/sec it did not significantly compromise the sensitivity (Figure S2 Supporting Information). In fact, at a speed rate of 30 scans/sec with the HTL at 450 °C, spectra at the higher range (*m/z* 700-900) still resulted in higher signal intensity when compared 150 °C at 10 scans/sec (Figure S3 Supporting Information). Moreover, spectra show consistency in peak shape with low to no peak splitting between 10 to 30 scans/sec, even for some lower intensity species (Figure S4 Supporting Information).

### Effect of DESI sprayer and HTL temperature on tissue histology

To evaluate the effect of the DESI-XS HP sprayer in combination with the HTL on tissue histology, H&E and PAS staining was performed on previously DESI-MSI scanned sections under different conditions. Data was acquired on three technical and three biological replicates scanned at 20 x 20 µm pixel size, at 10 scans/sec, with HTL set to 150 °C or 450 °C. The effect of solvent was assessed by keeping paired control sections, for 10 min, in a solution of the solvent mixture. A tissue section was placed inside the enclosed DESI source during the run of its paired section to evaluate the atmospheric temperature and moisture effect. A third control was kept at room temperature for the period of the DESI-MS run. After DESI scanning, sections were H&E stained and evaluated by an experienced kidney histopathologist (DJH) blinded to the conditions.

Post DESI-MSI analysis H&E and PAS-stained kidney sections (Figure 3) retain features of the renal cortex and medulla of the human kidney including glomeruli, proximal and distal tubules, collecting ducts, blood vessels, medullary rays and Bowman’s capsule.

**Figure 3.**
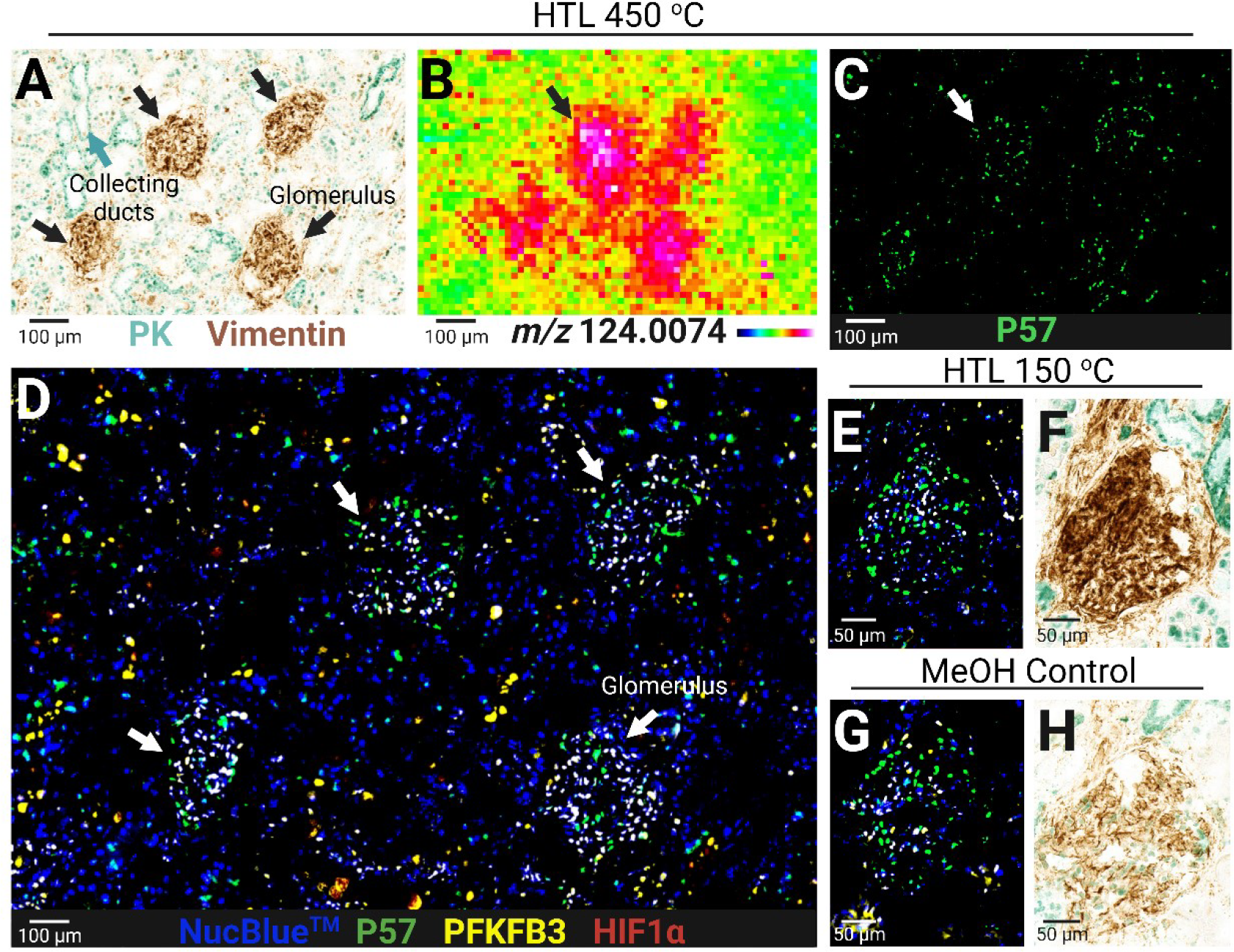
mIF and mIHC Post DESI-MSI. Protein epitopes and antibody binding for mIHC and mIF on kidney tissue remain intact post-DESI-MSI with the HTL at 150 °C and 450 °C. **(A)** mIHC image of collecting ducts (pan-cytokeratin^+^ (PK), green labelling) and glomeruli (vimentin^+^, brown labelling) post-DESI analysis with HTL at 450 °C, correlating to **(B)** higher taurine (*m/z* 124.0074) signal intensity and **(C)** podocytes (P57^+^) in glomeruli. **(D)** Composite of mIF-positive cells for P57 (podocytes, green labelling), PFKFB3 (glycolysis, yellow labelling) and HIF1α (hypoxia, red labelling) post-DESI with HTL 450 °C. **(E**,**G)** mIF and **(F**,**H)** mIHC image of the glomerulus of control and DESI analysed section at 150 °C. NucBlue™ was used as the nuclear marker.

**Figure 4.**
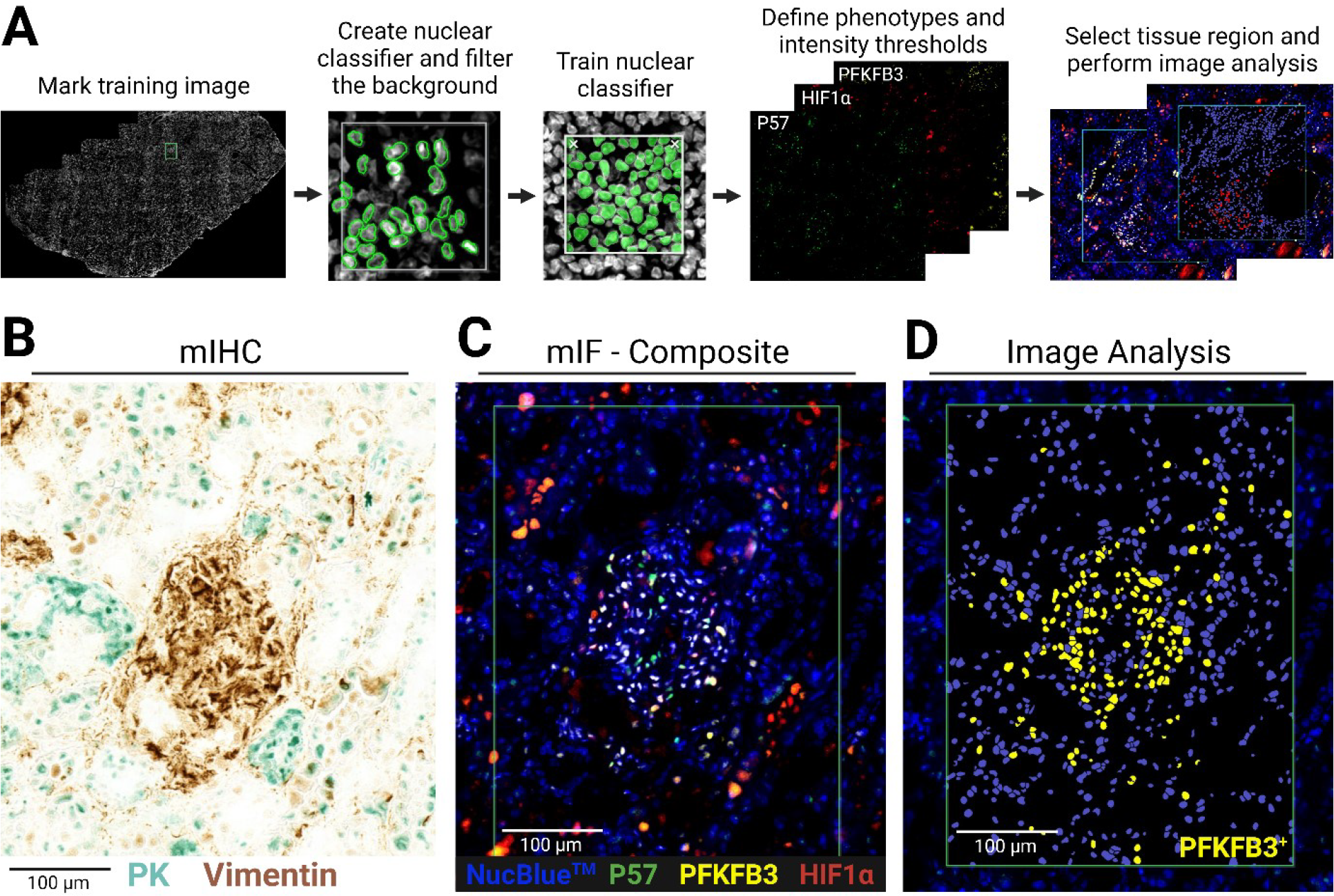
Image Analysis. (**A**) AI-enabled (HALO AI®) image analysis workflow of mIF data. **(B)** mIHC labelling of epithelial cells of the collecting ducts with pan-cytokeratin (PK, cyan chromogen) and glomerulus with vimentin and same region **(C)** mIF composite image of podocytes (P57^+^, green labelling), PFKFB3^+^ (yellow labelling) and HIF1α^+^ (red labelling) cells (NucBlue™ - Nuclear marker). **(C)** Nuclear segmentation and PFKFB3^+^ cells identified by AI-enabled image analysis. Tissue was analysed by DESI-MSI at 10 scans/sec with the HTL at 450 °C and H&E stained prior to mIF and mIHC.

Compared to the room temperature H&E controls, post-DESI sections and the MeOH controls show an intense eosin staining pattern with a more pronounced effect on the tissue submitted to analysis with the transfer line at 450 °C (Figure 3 and Figure S6 Supporting Information). The combination of MeOH and high temperature results in coagulation and fixation of proteins on the tissue, mainly in the intratubular spaces, whilst in the H&E control these proteins have likely been washed out during the staining process. The individual effect of the atmospheric temperature inside the DESI source and the MeOH:H^2^O mixture is confirmed by the controls that show a similar, however less pronounced, coagulation pattern. Even though fixed, proteins are detaching from the tubules and vessels’ walls resulting in loss of some fine morphology features remain recognizable. To assess, more in depth, the tissue integrity, PAS staining was used to highlight intact basement membranes of glomerular capillary loops and tubular epithelium across all DESI-MSI sampling conditions (Figure 3 and Figure S6 Supporting Information). As a HTL of 450 °C was determined optimal condition for lipid analysis but most impactful on the morphology of the tissue, the impact from the combination of faster scan speeds at the higher transfer line temperature was evaluated. DESI-MSI carried out at 20 and 30 scans/sec (Figure S7 Supporting Information) resulted in an identical protein coagulation and fixation pattern as seen previously at 10 scans/sec. However, increasing the sprayer speed reduced protein fragmentation, suggesting lower disturbance on tissue histology in these conditions when compared to the slower scanning rate.

### Multiplex IF and IHC achieved on the same tissue section post DESI-MSI and H&E

After confirming tissue integrity through histology post-DESI imaging experiments, we next examined the impact of the desorption and different HTL temperatures on tissue phenotyping. For that, a four-plex immunofluorescence and a double IHC labelling panel was performed on post-DESI sections and MeOH controls after they were stained for H&E and/or PAS. As the scanning rate of 10 scans/sec resulted in stronger morphological disturbance they were chosen for this assessment. A total of twelve sections were successfully stained for all four IF and two IHC markers after DESI-MSI analysis. For mIF, each phenotype signal was individually assessed and co-registered with the nuclear marker that confirmed the protein expression to their corresponding cellular structures (Figure 3). PFKFB3, HIF1α and P57 showed strong nuclei staining. P57 was positive for cells showing weaker NucBlue™ staining, and PFKFB3 was predominantly expressed in glomeruli cells and some cells of collecting ducts, confirmed by co-expression with pan-cytokeratin (PK) in mIHC images. Strong vimentin (brown) expression was observed in glomeruli and vascular structures whilst pan-cytokeratin (green) labelled epithelial cells of the collecting ducts. Increased taurine distribution in the glomerulus, observed in DESI-MS images, was confirmed by co-registration with mIHC (vimentin) and mIF (P57).

Epitope dilutions were optimized for optimal signal intensities and kept consistent between cases and different DESI imaging conditions (Table S1 Supporting Information). As previously confirmed by H&E on the same sections, 450 °C resulted in more protein fixation on the tissue leading to higher background signal in the mIF images. Immunohistochemical labelling was adequate for spatial data extraction after MSI, histology and mIF for samples subjected to both 150 °C and 450 °C DESI analysis. Both vimentin and PK showed different staining intensities between controls and DESI scanned sections. The fixation mechanism resultant from DESI at 150 °C appears to have a positive effect on IHC labelling by increasing the stain intensity. This effect was also seen in the sections that were PAS stained. Despite more severity of protein fixation and detachment at 450 °C, this did not negatively impact IHC staining as images from post-DESI-MSI and their MeOH control are comparable.

### Effect of DESI-MSI on image analysis by AI-enabled software

Following the pathologist’s evaluation, AI-enabled image analysis was used to identify and segment positive cells in the glomeruli for the three mIF markers. Figure 5 illustrates the workflow for image analysis where firstly, a nuclear segmentation classifier was built and trained. Subsequently, a threshold was set to each individual channel using nuclear staining (NucBlue™) as a guide to detect positive cells. As a result, image analysis accurately identified podocytes in the glomerulus and cells positive for PFKFB3 and HIF1α (Figure 5D) demonstrating segmentation and assignment can be performed on sections analysed at the most extreme conditions of 450 °C.

**Figure 5.**
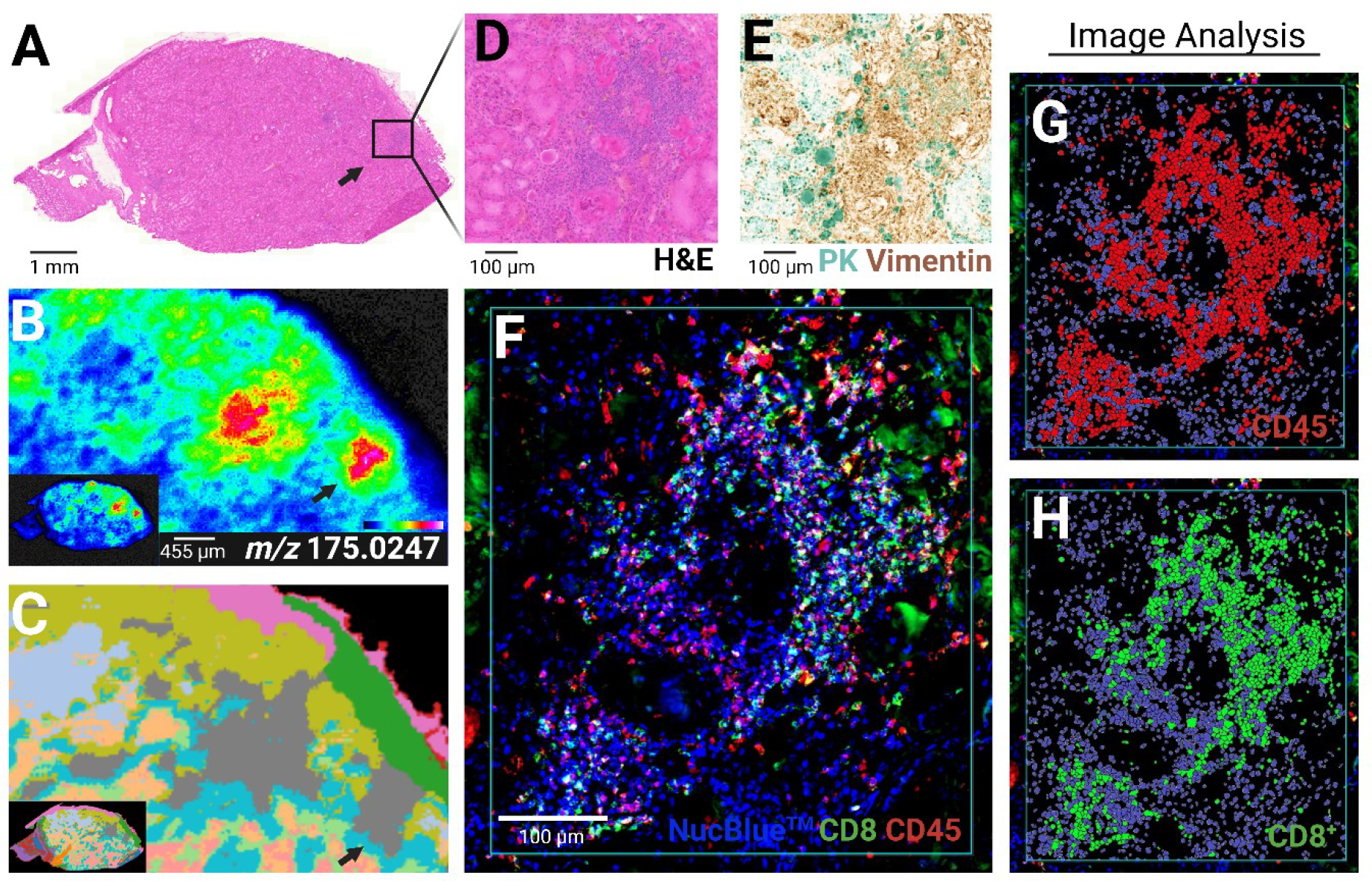
Differential metabolic fingerprints identified by DESI-MSI and UMAP segmentation confirmed immune infiltration by single-section multimodal imaging workflow. **(A)** Whole section H&E stain of a kidney tissue post-DESI scanning. **(B)** DESI-MS ion image of *m/z* 175.0247 (ascorbic acid) and **(C)** corresponding UMAP segmentation cluster of the MSI data. Detailed view of one of the areas with a higher concentration of ascorbic acid by **(D)** H&E showing fibrotic scars and immune cell infiltration; **(E)** mIHC labelling with pan-cytokeratin (PK, cyan labelling) and vimentin (brown labelling) expression indicating increased numbers of mesenchymal cells (fibroblasts) contributing to fibrosis and **(F)** composite image of mIF labelling of CD8^+^ (green) and CD45^+^ (red) cells. **(G**,**H)** Identified CD45^+^ and CD8^+^ cells through the application of HALO® AI image analysis to the mIF data. Section analysed with HTL at 450 °C.

### Application of the workflow: single-section DESI-MSI, H&E and mIF

DESI-MS images from one of the kidney sections previously had an unusual distribution for *m/z* 175.0247 (ascorbic acid). When subjected to a UMAP segmentation algorithm, 14 molecular clusters were identified. Of these, cluster 08, was found to spatially represent the area with increased ascorbic acid concentration. Re-evaluation of the H&E image revealed co-localisation of a higher intensity of ascorbic acid in regions of arteriosclerosis and hyalinisation of arterioles with focal chronic inflammatory cell infiltrate, consistent with chronic hypertension-related injury. To this section we then applied a different mIF panel and by digital image analysis identified CD45^+^ and CD8^+^ cells in these regions to demonstrate a real application of the evaluated workflow. Co-expression with vimentin indicates increased number of mesenchymal cells (fibroblasts) contributing to the fibrosis identified by the pathologist^15,16^.

## DISCUSSION

DESI-MSI allows for the direct analysis and visualisation of molecular information in tissue sections, providing insight into the spatial distribution of various compounds. DESI-MSI has successfully been applied for assessing lipid metabolism in different pathologies such as diabetic kidney disease^17^, breast and thyroid cancer^18^, clear cell renal cell carcinoma^9^ and brain tumours^19^. Here we have shown that increasing the temperature of the HTL results in greater sensitivity of lipid species in the range of *m/z* 600-1000, however, this is not a global metabolite effect, as the signal of most small molecule species (*m/z* 50-350) reduced with increasing HTL temperature. However, some compounds, such as taurine, increased. We have not explored in depth the effect of temperature on small molecules; however, small molecules have been shown to be thermally unstable^20^. This increase in sensitivity of small molecules at higher HTL temperature may be artefactual because of in-source fragmentation, ion competition, desorption degradation due to the high temperature of the HTL or on-tissue degradation, as tissues sat in the DESI source under a heated atmosphere for several hours during analysis. Therefore, care should be taken when interpretating data from samples submitted to heated conditions.

Integration of DESI-MSI into pathology workflows offers the potential to complement traditional histological techniques and enhance our understanding of disease processes. Here we explored the capacities of the redesigned DESI sprayer and HTL at different temperatures and its integration with multiple histology, mIF, mIHC and image analysis for full histological assessment from a single section. Thus, allowing direct correlation of metabolites and protein distribution with histological features from patient-derived human tissue.

Long acquisition time is one of the limitations of MSI technologies with samples taking up to several hours to be analysed. Patient-derived biopsies, however, are usually small pieces of tissue that generally require less analysis time. In addition, changing MSI acquisition parameters helps decrease the time for data acquisition. Analysts, nonetheless, must be aware that these changes might mean sacrifice sensitivity and spatial resolution. By using the novel DESI XS sprayer in conjunction with the HTL and increasing scan speeds from 10 scans/sec to 30 scans/sec we have obtained MSI results almost 3-fold faster with no significant loss of sensitivity and resolution. Furthermore, sacrificing sensitivity for faster runs when using the HTL still resulted in better lipid signal than at slow acquisition rate at lower temperature. This, for small areas of tissue, such as patient biopsies, could offer runs of just a few minutes with high quality data.

In biomedical applications, correlation between the chemical information, obtained by DESI-MSI, and histology allows for correct assignment of ion images to their corresponding anatomic structures. Whilst the use of adjacent sections in some cases might still be sufficient for structure assignment, protein expression greatly varies from one section to other, not truly corresponding to the chemical results obtained by MS. In this work various DESI-MS imaging conditions have been explored to assess the effects on tissue and their impact on downstream processes. The results show that indeed DESI-MSI, whilst a soft ionisation technology, still does create alterations to the tissue. H&E and PAS, as standard stains used in kidney biopsy, showed that both temperature^25^ and solvent are responsible for the protein coagulation and detachment from the kidney tubular structures. And whilst H&E of 450 °C samples was found to highlight this effect, PAS stain showed no significant difference between all the different DESI conditions in comparison to the controls.

The speed of the sprayer across the surface of the sample was also shown to slightly impact the morphology. The harshest effect seen at 10 scan/sec in comparison to 30 scans/sec agrees with the increased sensitivity observed in the MS results. The slower movement of the sprayer enhances the extraction of the sample components into the solvent increasing the intensity of the signal, but also results in a stronger physical and chemical effect on the cellular and acellular structures of the tissue. Although there is a loss in fine morphology observed in all sampled conditions, this loss has minimal effect on interpretation. mIF and/or mIHC post-MSI analysis is strongly advantageous to obtain a rich correlation between protein expression and phenotypic features with metabolic markers. In this work we have shown that mIF and IHC can be performed on sections that have been analysed by DESI and H&E stained. Although, intensity differences were seen for both mIF and IHC markers between samples submitted to DESI under different analysis conditions and serial non-DESI scanned sections. These differences might be a result of the DESI analysis but are also related to the pre-analytical steps. Though we have optimized the cryosection step to obtain same thickness sections, variability is still present and is operator dependent. This is an important step in MSI and histochemistry and must be carefully considered, as thicker sections can result in stronger shattering. Furthermore, tissue thickness is known to positively affect the fluorescence intensity^26^ and with the sprayer and high temperature, higher intratubular protein coagulation may result in an increase of background signal. However, we have demonstrated that with extreme DESI-MSI conditions and pre-analytical variables, AI segmentation and phenotyping was successful in both mIF panels we applied.

## CONCLUSION

A workflow incorporating Mass Spectrometry Imaging (MSI) for co-registration of metabolic changes with molecular markers aided by machine learning was developed, evaluated, and applied to a single section of non-malignant human kidney demonstrating the potential for DESI-MSI to secure its position as a crucial tool in next – generation pathology.

## ACKNOWLEDGEMENTS

This work was supported by The Melville Trust (G.M.Z) and the KATY project (IU, DJH) that has received funding from the European Union’s Horizon 2020 research and innovation programme under grant agreement No. 101017453, and in part by the Industrial Centre for AI Research in Digital Diagnostics (iCAIRD) which is funded by Innovate UK on behalf of UK Research and Innovation (UKRI) (project number 104690). A.L.D and D.J.H are also funded by NuCana plc.

## Notes

### Competing Interest Statement

The authors have declared no competing interest.

