## Supplementary Information for "Spatial Metabolomics co-registered with Multiplex Phenotyping for the Evaluation of Human Kidney Tissue"

**Supplementary Methods**

Table 1. Antibody characteristics and dilutions

| Methodology | Antibody | Company | Product code | Dilution |
|---|---|---|---|---|
| mIHC | Vimentin | Cell signalling tech | 5741 | 1:50 |
| | Pan-cytokeratin | Dako | Z0622 | 1:50 |
| mIF | P57 | Santacruz | sc56341 | 1:100 |
| | HIF1α | Abcam | ab51608 | 1:600 |
| | PFKFB3 | Abcam | ab181861 | 1:200 |
| | CD45 | Abcam | ab40763 | 1:500 |
| | CD8 | Agilent | M710301-2 | 1:400 |

**Single-section histochemical staining, multiplex immunofluorescence and multiplex immunohistochemical labelling protocol.**

After each DESI experiment, tissues were kept in 10% Tween® 20 detergent (TBST) buffer for 5 min to remove hydrogel, immersed in haematoxylin for 3 min, then washed in three consecutive water baths and immersed in TBST buffer for 1 min. Slides were then immersed in eosin for 10 sec, followed by consecutive washes in ethanol at 50, 80 and 100% baths. For dehydration, slides were immersed for 5 min in three different 100% xylene baths. Samples were then fixed with DPX glue and a coverslip.

After H&E stained, selected sections were immersed in xylene until coverslip and all DPX glue was removed. Tissues followed a rehydration step of 2 min in 100, 80 and 50% ethanol and 2 min in distilled water. Slides were kept flat; sections were covered with 1% periodic acid for 5 min and then washed in tap and distilled water prior to being covered with Schiff's reagent (1:4) for 10 min. Then sections were kept under warm running tap water for 5 min and counterstained with haematoxylin for 3 min following the same sequence above mentioned for H&E staining (skipping the eosin step) and fixed with a coverslip.

Following H&E or PAS sections were uncovered in xylene solution and kept in TBST prior to mIF staining.

**Supplementary Results**

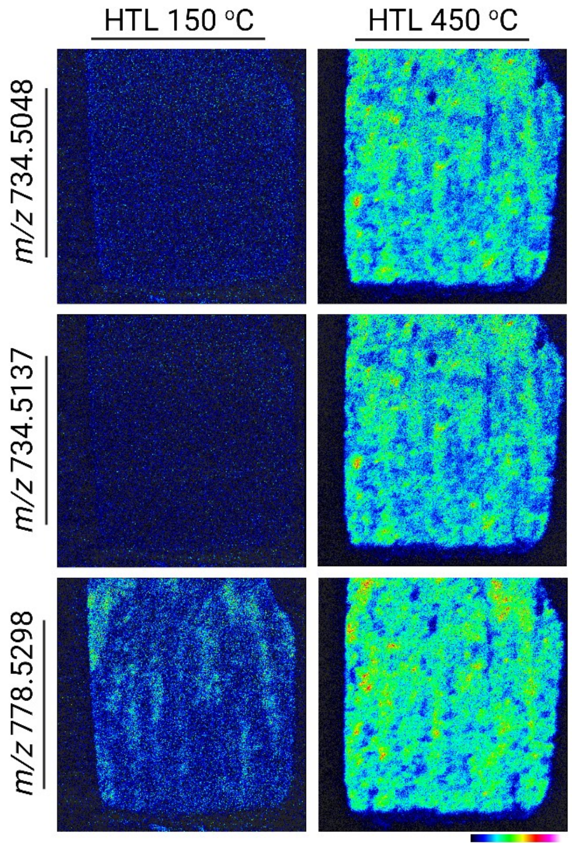


Supplementary Figure S1. Ion images of *m/z* values with increased sensitivity when analysed with the DESI XS HTL at 450 °C.

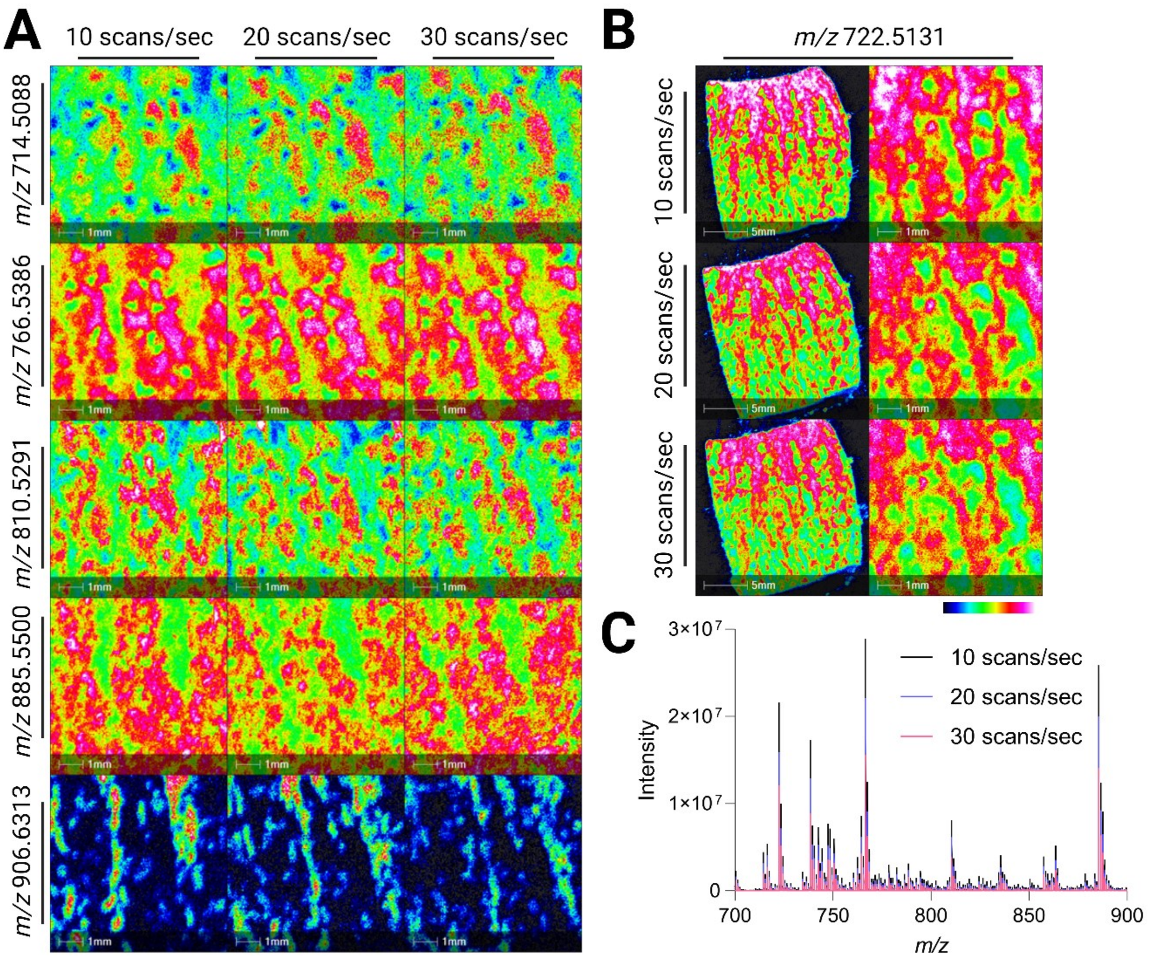


Supplementary Figure S2. **(A)** Ion images of lipids analysed by DESI-MSI with the HTL at 450 ºC at different scan speeds, *m/z* 714.5088 (PE 34:2), *m/z* 766.5386 (PE 38:4), *m/z* 810.5291 (PS 38:4), *m/z* 885.5500 (PI 38:4) and *m/z* 906.6313 (SHexCer 42:1;O3). **(B)** Whole human kidney tissue section and zoomed-in region ion images of *m/z* 722.5131 (PE O-36:5) analysed by DESI-MSI with the HTL at 450 ºC at different scan rates. (C) DESI-MSI spectra of kidney tissue sections in the range *m/z* 700-900 showing a decrease in sensitivity when increasing the acquisition speed when analysed with the HTL at 450 ºC.

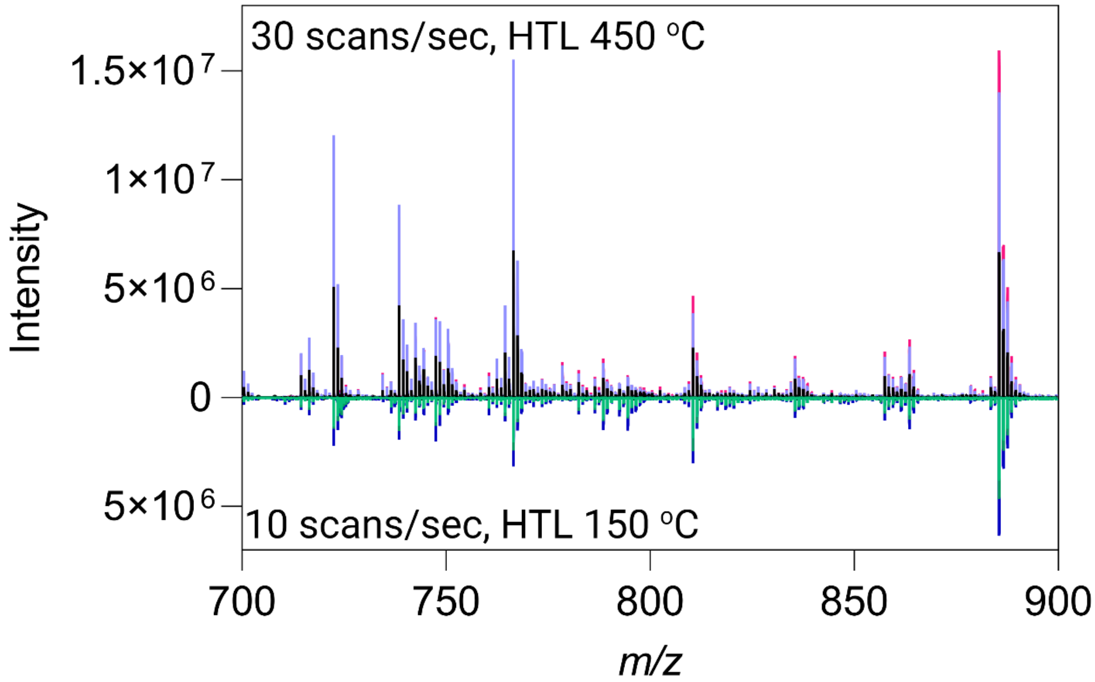


Supplementary Figure S3. **DESI-MS spectra of kidney tissue obtained with HTL set to 450 ºC and scan speed of 30 scans/sec (pink, purple and black spectra), and at 150 ºC and 10 scans/sec (blue, light and dark green spectra)**. We used data from the previous experiment for the transfer line temperature at 150 ºC for this comparison. Same size ROI were drawn on the same region of all 6 sections. As sections are not consecutive but are from the same tissue, we compared the spectra of sections scanned at 10 scans/sec at 450 ºC for the transfer line experiment with the spectra of the replicates of the 10 scans/sec scan speed experiments (data not shown). As intensities were consistent between both experiments, we concluded that a comparison between 150 ºC at 10 scans/sec and 450 ºC at 30 scans/sec was appropriate.

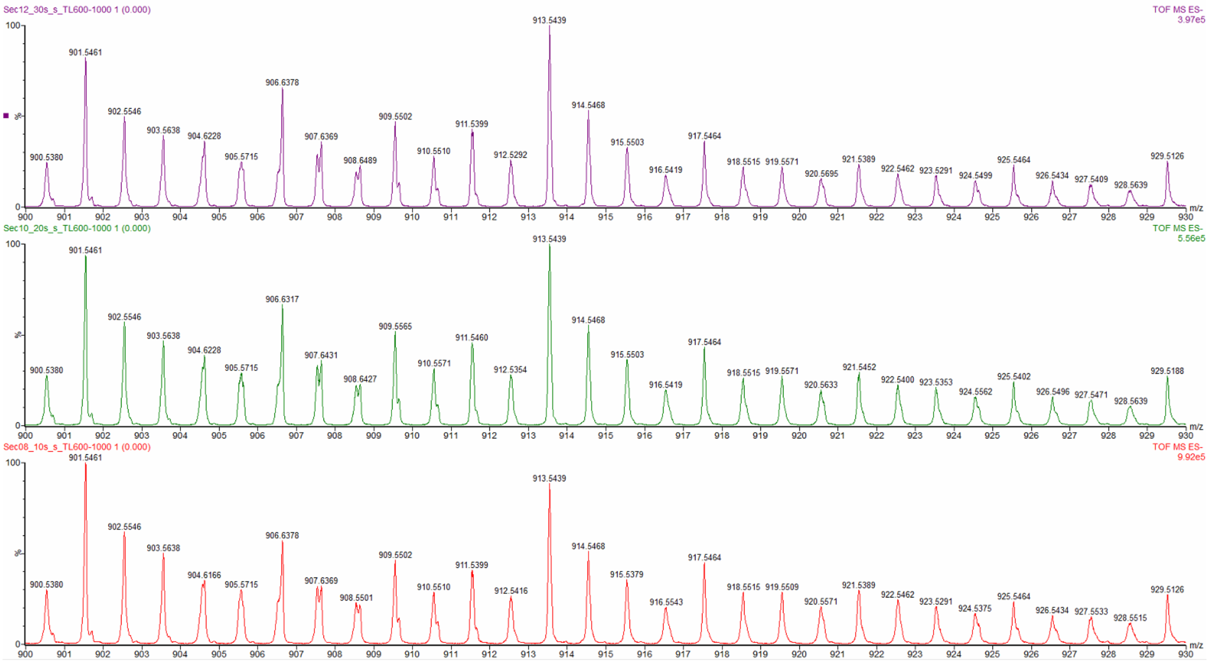


Supplementary Figure S4. **Illustration of peak shape consistency for low-intensity peaks at different scan speeds in the range *m/z* 900-930**. Red spectra show data acquired at 10 scans/sec, green spectra at 20 scans/sec and purple spectra at 30 scans/sec.

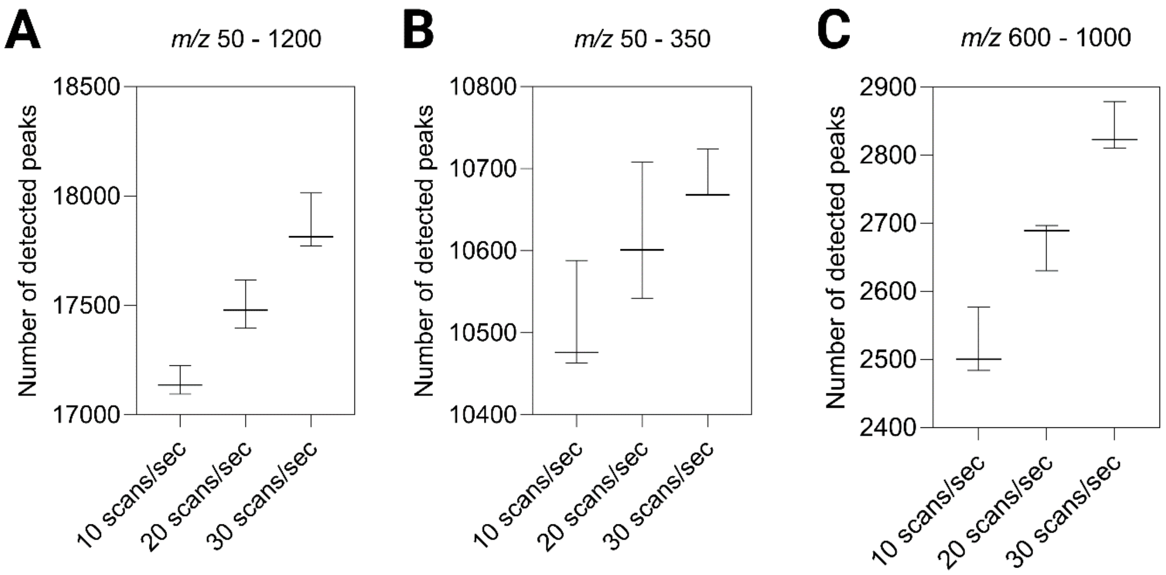


Supplementary Figure S5. **Number of detected features at different DESI-MSI scan speeds. (A)** Number of peaks detected in the entire acquired range, *m/z* 50 – 1200, **(B)** in the low molecular weight range, *m/z* 50-350, and **(C)** in the high molecular weight range, *m/z* 600-1000. (n=3)

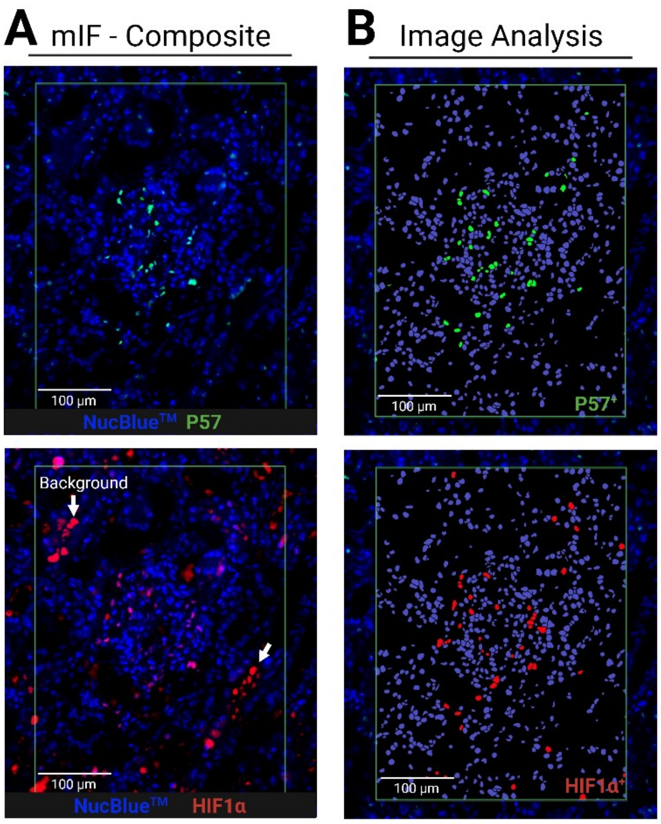


Supplementary Figure S6. **(A)** mIF images of nuclear marker (NucBlue™), podocytes (P57$^+$, green labelling) and HIF1α$^+$ cells (red labelling) of the same region of a tissue analysed by DESI-MSI with the HTL at 450 °C at 10 scans/sec. **(B)** AI-enabled nuclear segmentation and identification of cells positive for P57 and HIF1α.
